# Combination of memantine and alpha7 nicotinic acetylcholine receptor ligands exerts superior efficacy over monotreatments to improve cognitive performance of aged rats

**DOI:** 10.1101/2023.09.20.558650

**Authors:** Nóra Bruszt, Zsolt Kristóf Bali, Lili Veronika Nagy, Kornélia Bodó, Péter Engelmann, István Ledneczki, Zsolt Némethy, Balázs Lendvai, István Hernádi

**Affiliations:** Translational Neuroscience Research Group, Centre for Neuroscience, Szentágothai Research Centre, University of Pécs, 20 Ifjúság str., H-7624 Pécs, HUNGARY; Institute of Physiology, Medical School, University of Pécs, 12 Szigeti str., H-7624 Pécs, HUNGARY; Grastyán Endre Translational Research Centre, University of Pécs, 6 Ifjúság str., H-7624 Pécs, HUNGARY; Department of Neurobiology, Institute of Biology, University of Pécs, 6 Ifjúság str., H-7624 Pécs, HUNGARY; Institute of Immunology and Biotechnology, Medical School, University of Pécs, 12 Szigeti str., H-7624 Pécs, HUNGARY; Department of Chemistry, Gedeon Richter Plc., 19-21 Gyömrői str., H-1103 Budapest, HUNGARY; Pharmacology and Drug Safety Research, Gedeon Richter Plc., 19-21 Gyömrői str., H-1103 Budapest, HUNGARY; Department of Richter, Semmelweis University, 26 Üllői str. H1085 Budapest,. HUNGARY

## Abstract

Combination treatments based on pharmacological interactions at α7 nAChRs are promising therapeutic approaches for neurocognitive disorders (NCDs). Memantine, an already approved medication in some NCDs, may not only act as an antagonist of the glutamatergic NMDA receptors (NMDAR) but also serve as an antagonist of the cholinergic α7 nAChRs. Here we set out to utilize a combination treatment regime with an α7 nAChR-selective agonist (PHA-543613) and a novel proprietary α7 nAChR ligand with marked positive allosteric modulator (PAM) activity (CompoundX). The cognitive efficacy of combination treatments was tested in a naturally aged rat model. Naturally aged rats showed marked cognitive decline in the novel object recognition (NOR) test, and they displayed pathological changes at the molecular level in terms of various inflammatory markers. In addition, aged rats also exhibited cholinergic changes such as mRNA upregulation of α7 nAChRs. Memantine-PHA-543613 and memantine-CompoundX combination treatments successfully alleviated the age-related decline of recognition memory of rats by exceeding the null-effects of the corresponding subtherapeutic levels of monotreatments. Results indicate a positive interaction between memantine and α7-nAChR agonists and PAMs, which reflects a prominent role of α7 nAChRs in the cognitive enhancement of combination treatments especially in age-related cognitive decline. Moreover, the putative direct action of memantine on α7 nAChRs may also contribute to the observed synergistic interaction between memantine and other selective α7 nAChR compounds.

## 1. Introduction

Aging is generally associated with emerging neurocognitive disorders (NCDs), that pose serious public health challenges worldwide (Hou et al. 2019; Llaurador-Coll et al. 2023). These age-related disorders are characterized by progressive neuronal dysfunction and deterioration of cognitive abilities and activities of daily living. Multiple cellular and molecular changes can be associated with age-related cognitive decline in humans and in preclinical animal models, and neuroinflammation is a general process underlying these pathological changes (Ownby 2010; Simen et al. 2011; Shobin et al. 2017; Angelova and Brown 2019; Thadathil et al. 2021). Thus, promising future treatment strategies should address neuroinflammatory mechanisms underlying the neurophysiological dysfunctions of the aging brain.

Currently, the non-competitive NMDA receptor antagonist memantine and different cholinesterase inhibitors (e.g., donepezil) can be offered as therapeutic options for patients with NCDs, however, several clinical observations indicate that these medications may provide only moderate and transient symptomatic benefits with very limited or no disease-modifying potential (Raina et al. 2008; Tsoi et al. 2016; Balázs et al. 2022). Therefore, development of new treatment strategies is still an unmet medical need and the discovery of new avenues for the treatment of NCDs would be crucial in the field (Sharma et al. 2019). The alpha7 nicotinic acetylcholine receptor (α7 nAChR) is considered a promising target to treat NCDs because it is primarily involved in learning and memory processes (Wallace and Bertrand 2015; Corradi and Bouzat 2016) and also plays a role in immune regulation (Zdanowski et al. 2015). It is well known that signaling through α7 nAChRs enhances both cholinergic and glutamatergic transmission contributing to memory improvement in preclinical disease models (Bali et al. 2015, 2017, 2019b). In addition, in recent years, stimulation of α7 nAChRs expressed by glial cells has been shown to reduce neuroinflammation by activating the cholinergic anti-inflammatory pathway (Conejero-Goldberg et al. 2008; Egea et al. 2015; Corradi and Bouzat 2016; Maurer and Williams 2017; Mizrachi et al. 2021) and, thus, also contributing to the reported therapeutic efficacy of α7 nAChRs in preclinical models of NCD (Echeverria et al. 2016; Foucault-Fruchard et al. 2017).

Combination therapies may be highly promising novel approaches in the treatment of age-related NCDs, since the co-application of pharmacological agents with different mechanisms of action may result in a superadditive (synergistic) increase in efficacy and/or enables the use of lower doses, while achieving the same effectiveness with less possible side-effects. Furthermore, combination treatments may offer a complex influence on the dysfunction of signaling pathways and pathological processes involved in NCDs (Parsons et al. 2013), which may add up in beneficial disease-modifying pharmacological effects. Although the combination of acetylcholine esterase inhibitor (AChEI) donepezil with glutamatergic antagonist memantine is already in use in the treatment of Alzheimer’s disease, clinical evidence is ambiguous about its superior cognitive outcomes over the corresponding monotherapies (Deardorff and Grossberg 2016; Knorz and Quante 2021). Consistent with this, preclinical studies also confirmed that combination treatment with AChEIs and memantine provides little or no beneficial outcome in different behavioral tests (e.g. radial arm maze, classical eyeblink conditioning) in animal models (Wise and Lichtman 2007; Woodruff-Pak et al. 2007). In contrast, available results suggest a more promising outcome in cases when memantine is combined with compounds acting directly on the α7 nAChRs (Koola et al. 2018). For example, co-administration of memantine with galantamine (which acts not only as an AChEI but also as a positive allosteric modulator (PAM) of α7 nAChRs) alleviates scopolamine-induced memory deficits in mice (Busquet et al. 2012) and delays natural forgetting in rats (Nikiforuk et al. 2016) with higher efficacy compared to the corresponding monotreatments. These findings are further supported by two recent studies from our laboratory, in which the combined effect of memantine and the selective α7 nAChR agonist PHA-543613 was tested on different cognitive functions in a scopolamine-induced transient amnesia model. First, we reported, that rats receiving memantine-PHA-543613 treatment showed better working memory performance in the spontaneous alternation task compared to rats receiving monotreatments (Bali et al. 2019a). Furthermore, our most recent findings indicate that the same treatment combination improved both short-term memory and recall of long-term memory of rats in the Morris water maze test (Bruszt et al. 2021). Based on the above findings, we previously suggested a potential pharmacodynamic interaction between α7 nAChR ligands and memantine, which was also supported by previous studies that reported higher affinity of memantine to α7 nAChRs than to NMDA receptors (Aracava et al. 2005a; Banerjee et al. 2005).

Taken together, the aim of the current study was to further investigate the behavioral level interaction between memantine and two different α7 nAChR agents (the orthosteric agonist PHA-543613 and a novel compound with PAM characteristics: CompoundX, a proprietary compound of Gedeon Richter Plc., Hungary, described in the patent WO 2020/012423 A1) in a preclinical animal model of age-related cognitive deficit. In the present study, naturally aged rats were used as they show age-related cognitive impairment and certain emerging pathological processes including neuroinflammation and apoptotic cell death, posing as one of the most relevant models of spontaneous, age-related cognitive decline in humans (Maher et al. 2005; Frank et al. 2006). Since aging is not a uniform process neither in rodents nor in humans, individuals of the same age may exhibit varying cognitive performance in behavioral tasks which may mask smaller drug-induced effects. To tackle the above problem of the model, our further aim was to assess key molecular-level indicators of pathological and healthy aging animals by investigating the mRNA and protein expression of inflammatory biomarkers (IL-1β, MIP-1α), ciliary neurotrophic factor (CNTF), and α7 nAChR in both memory-impaired and unimpaired animals.

## 2. Materials and Methods

### 2.1. Animals

Altogether 65 (comprising of 12-15 subjects/group) naturally aged (28 months old) and 12 young (4 months old) male Long Evans rats (Charles River Laboratories, Calco, Italy) weighing 350–500 g were applied in the current study. Animals were pair-housed under 12/12 h daily light/dark cycle with controlled temperature and humidity in the animal house of the Szentágothai Research Centre, University of Pécs, Hungary. In the animal house, the lights were ON from 7 a.m. to 7 p.m., and the animals were tested in the light period. Rats were fed daily with controlled amount of food (17 g/animal laboratory chow) to prevent the development of obesity and other related health problems. Water was available ad libitum. The experiments were approved by the Animal Welfare Committee of the University of Pécs, and the National Scientific Ethical Committee on Animal Experimentation (ÁTET) at the Ministry of Agriculture (license no.: BA02/2000–25/2015, BA02/2000-30/2021). All procedures fully complied with the Decree No. 40/2013. (II. 14.) of the Hungarian Government and the EU Directive 2010/63/EU on the Protection of animals used for scientific purposes.

### 2.2. Behavioural assessment

#### 2.2.1. Novel object recognition test

Long-term declarative memory of the animals was evaluated with the NOR test paradigm. The experiments were carried out in an open field box which was made of grey-colored plywood, in size of 57.5 x 57.5 cm (length x width) surrounded by 39.5 cm high walls. The test protocol consisted of two trials: an acquisition trial followed by a test trial after 24 hours retention interval. In the acquisition trial, two identical objects were placed near the left and right corners of the box, and the rats were allowed to explore the environment and the objects for 3 min. For the retention interval, animals were returned to their home cages and were transferred to the animal house. After 24 hours, novel object recognition was assessed in the test trial. One of the identical objects was changed with a novel one and rats were allowed to explore the objects for 3 min. In a within-subject experimental arrangement, four different object pairs were used that were distributed randomly between animals and experimental sessions in a counterbalanced Latin-square design.

Because of the innate novelty-seeking behavior of rats, successful recognition of the familiar object (normal memory function) is supposed to be manifested in the preference to explore the novel object. Thus, the time spent with familiar (Ef) and novel (En) objects was recorded, and a discrimination index (DI) was calculated based on the following formula:

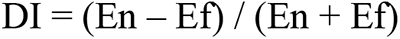

The DI appeared to be a positive number if the subject explored the novel object for a longer duration compared to the familiar object, the DI was negative if the familiar object was observed longer, and the DI was around zero if the two objects were observed for an almost equally long duration. Rats who did not observe the two objects together for at least 5 sec or observed only one of the objects (i.e., DI=±1.00) were excluded from the analysis of the given session.

#### 2.2.2. Drugs and routes of administration

PHA-543613 hydrochloride (Cat. No. 3092, Tocris), and memantine hydrochloride (Cat. No. 0773, Tocris) were dissolved in physiological saline and were applied in a final injection volume of 1 ml/kg. CompoundX (a proprietary compound of Gedeon Richter Plc., Hungary, described in the patent WO 2020/012423 A1, synthesized by Gedeon Richter Plc.) was dissolved in 5% Tween (in PBS) and was applied in a final injection volume of 2 ml/kg. All compounds were injected subcutaneously (SC) 45 min before the acquisition trial. In co-administration treatments, memantine and PHA-543613 or memantine and CompoundX were administered consecutively in two separate subcutaneous injections. When mono-treatments were tested, drugs that were not administered were replaced with the corresponding vehicle.

#### 2.2.3. Experimental design

In the first part of the study, cognitive behavior of the animals was measured using the NOR paradigm. First, the baseline performance of aged and young rats was compared. After the successful demonstration of cognitive impairment in aged rats, pharmacological effects of monotreatments with memantine, PHA-543613, or CompoundX, as well as co-administrations of memantine and one of the cholinergic agents, were tested in aged animals. In the pharmacological experiments, treatments were applied in a counterbalanced, within-subject design to achieve a fully randomized sequence of different treatments. Monotreatments with memantine were applied in the following doses: 0.1 mg/kg, 0,3 mg/kg, and 1.0 mg/kg (Mem0.1, Mem0.3, and Mem1.0, respectively). PHA-543613 was administered in doses of 0.3 mg/kg, 1.0 mg/kg, and 3.0 mg/kg (PHA0.3, PHA1.0, and PHA3.0, respectively) and CompoundX was administered in the doses of 0.3 mg/kg, 1.0 mg/kg, and 3.0 mg/kg (CPDX0.3, CPDX1.0, and CPDX3.0, respectively). Then, co-administration of subeffective memantine and PHA-543613 doses were tested against the effects of the corresponding monotreatments by applying the following treatments in a counterbalanced Latin-square design: vehicle alone (VEH), memantine monotreatment in 0.01 mg/kg dose (Mem0.01), PHA-543613 monotreatment in 0.1 mg/kg dose (PHA0.1), and co-administration treatment (Mem0.01&PHA0.1). Similarly, the efficacy of memantine-CompoundX co-administration in comparison to the monotreatments was tested in a separate experiment using the following treatments: vehicle alone (VEH), memantine monotreatment in 0.01 mg/kg dose (Mem0.01), CompoundX monotreatment in 0.1 mg/kg dose (CPDX0.1), and co-administration treatment (Mem0.01&CPDX0.1).

### 2.3. Brain tissue collection and preparation

After completing the behavioral experiments, animals were anesthetized with an overdose of pentobarbital and were transcardially perfused. A subgroup of young and aged rats was perfused with 0.9% saline, their brains were rapidly removed and dissected into left and right neocortex (CTX), striatum (STR), and hippocampus (HC). The brain samples were frozen immediately in liquid nitrogen and stored at -80 °C until biochemical analyses were performed.

### 2.4. RNA isolation, cDNA synthesis, and quantitative real-time PCR

The total RNA of the brain samples was extracted according to the manufacturer’s protocol by applying the NucleoSpin RNA kit (Macherey-Nagel, Düren, Germany). The quality and quantity of RNA were assessed at 260 nm using a NanoDrop 1000 spectrophotometer (Thermo Fisher Scientific, Waltham, USA). The cDNA was constructed from total RNA with High Capacity cDNA reverse transcription kit (Thermo Fisher Scientific) in 20 μl reactions using random hexamers following the manufacturer’s protocol. The resulting cDNA was stored at −20 °C. Target gene expressions were measured using real-time PCR using Maxima SYBRGreen MasterMix (Applied Biosystems, Waltham, USA) with an ABI Prism 7500 instrument (Applied Biosystems). The cDNAs were applied as a template for the amplification reactions. All samples were tested in duplicates. Primers were designed by Primer Express software (Thermo Fisher Scientific) considering the exon-intron boundaries for all target genes (Table 1). Cyclophilin A (CycA) was used as a housekeeping gene for the quantification of RNA. The thermal profile started at 95 °C for 10 min, 40 cycles of 35 sec at 95 °C, 35 sec at 60 °C, and 1 min at 72 °C.

**Table 1.**
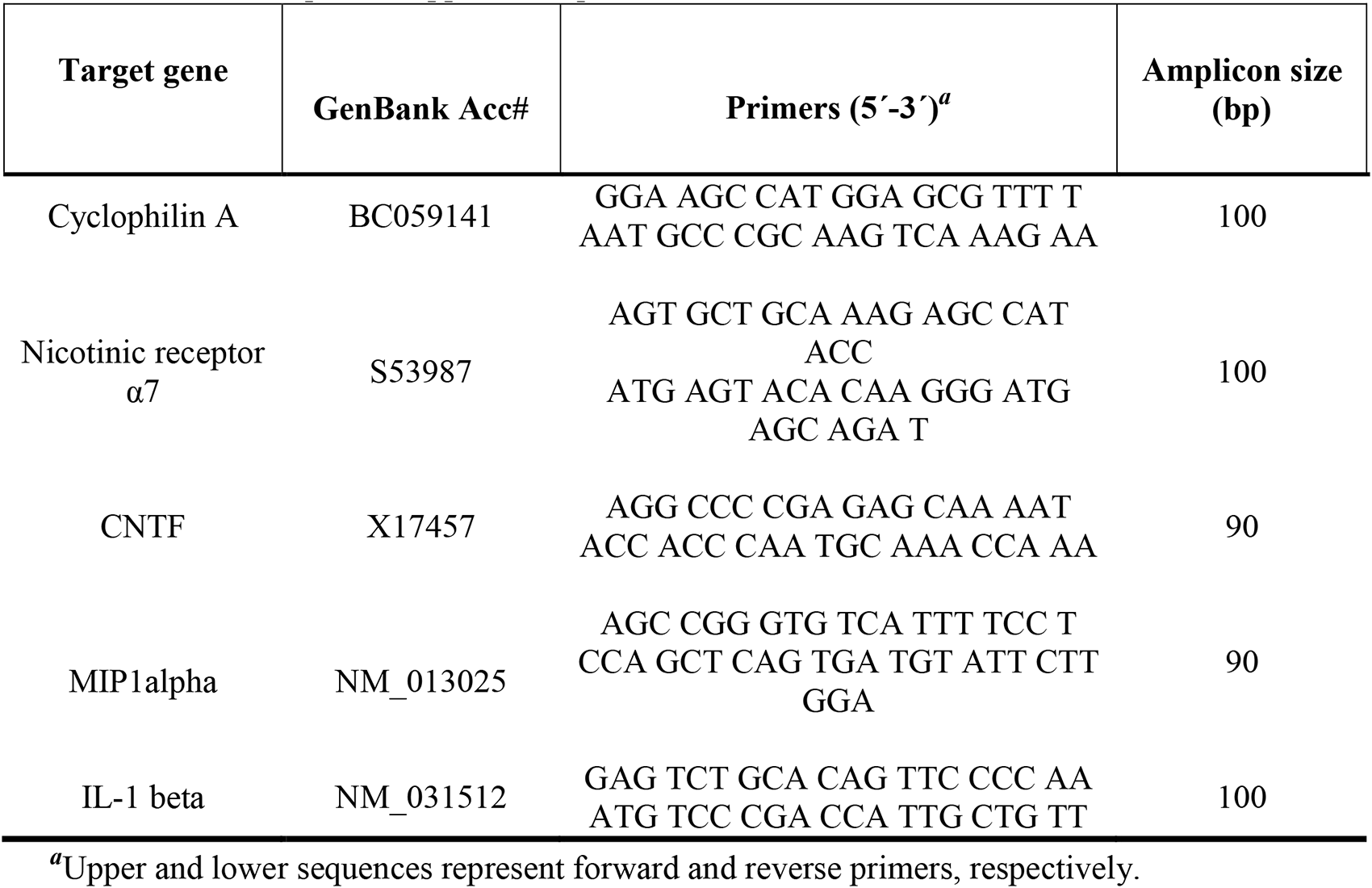
Primer sequences applied for qPCR.

### 2.5. Protein isolation and ELISA test

Total protein from the different brain regions was isolated in RIPA lysis buffer (50 mM Tris– HCl, pH: 8.0, 150 mM NaCl, 1% NP-40, 0.5% Na-deoxycholate, 5 mM EDTA, 0.1% SDS). The total protein concentrations of the samples were measured using BCA Protein Assay kit (Pierce, Rockford, IL). Quantification of IL-1β, MIP-1α, CNTF, and α7 nAChR was carried out using Abbexa (Cambridge, UK) ELISA kits with the following catalog numbers, respectively: abx155713, abx155822, abx155360, abx556026. ELISAs were performed according to the manufacturer’s protocols. Immediately after developing the final color-reaction of the assay, the absorbance was measured at 450 nm with an iEMS Reader microplate reader (Thermo Fisher Scientific).

### 2.7. Statistical analysis

Data were expressed as mean ± standard error of the mean (SEM). Statistical analyses were performed using the IBM SPSS 20.0 software. In the NOR test, the time spent exploring the familiar and novel objects was compared using paired samples T-test to assess whether rats after a given treatment show normal recognition memory performance. The DIs of aged and young animals were compared using independent samples T-test. In pharmacological experiments, the main effects of treatments on DI were analyzed using linear mixed-effect model. Following a significant main effect, effects of the tested treatments were compared with the effect of the vehicle using post-hoc LSD test. RNA and protein expression levels of young, memory impaired, and unimpaired aged groups were compared using univariate ANOVA and post-hoc LSD test.

## 3. Results

### 3.1. Effects of aging on long-term recognition memory performance

To determine age-related cognitive impairment, novel object recognition memory of aged and young animals was assessed and compared. In the test trial after 24 hours retention interval, young animals spent significantly more time exploring the novel object than the familiar one (exploration time of the novel vs. familiar object: 9.8**±**0.9 vs. 4.6**±**0.4, t =6.666, df = 11, p < 0.001). As expected, no discrimination between the familiar and the novel objects was observed in the case of aged rats (novel vs. familiar: 8.1**±**1.0 vs. 6.1**±**0.7, t =1.526, df = 10, p = 0.158) (Fig.1A). In addition, the DI of aged rats was significantly lower compared to young rats (aged vs. young: 0.130**±**0.085 vs. 0.352**±**0.039, t =2.426, df = 21, p = 0.024) (Fig. 1B). Results indicate that aged rats show remarkable deficits in the long-term declarative (recognition) memory.

**Figure 1.**
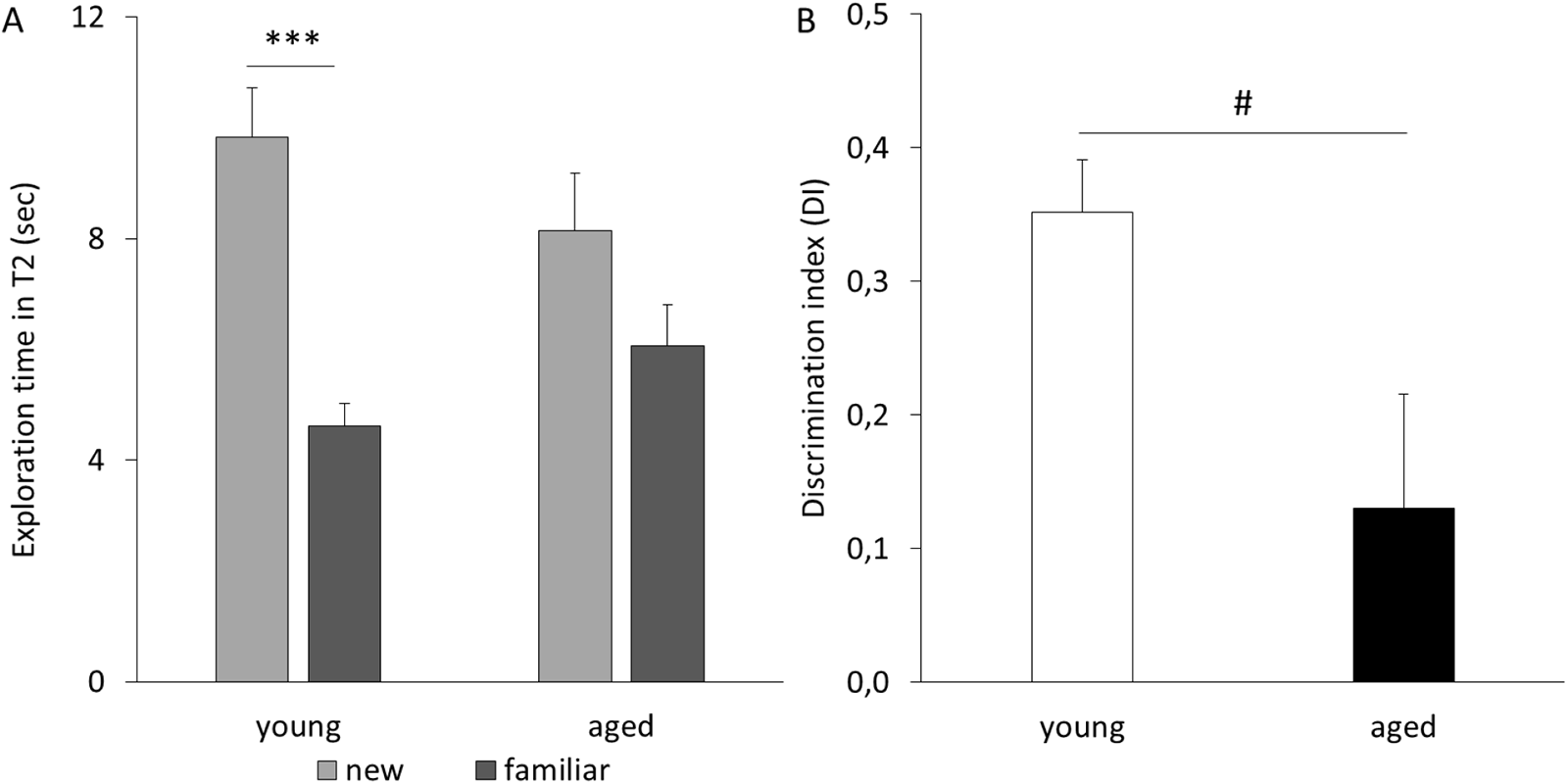
Comparison of aged and young animals in the NOR test. A: Young rats spent significantly more time exploring the novel object compared to the familiar one, while aged rats did not discriminate between the novel and the familiar objects. B: DI of aged rats was significantly lower compared to young rats. Asterisks mark significant difference in the time spent with the exploration of the novel and the familiar objects: ***p<0.001 (paired samples T-test). Hash indicates significant difference between the DI of aged and young animals: #p<0.05 (independent samples T-test).

### 3.2. Effects of monotreatments with cognitive enhancers memantine, PHA-543613, and CompoundX

The dose-response relationship of procognitive effects of memantine and the α7 nAChR ligands PHA-543613 and CompoundX was studied as the modulation of natural memory decline caused by aging. Unlike vehicle treatment, the administration of memantine (0.1 and 1.0 mg/kg) significantly increased the time spent with the exploration of the novel object as compared to the familiar object (time spent observing the novel vs. familiar: VEH: 9.3±1.6 vs. 7.3±1.1, t=1.489, df=10, p=0.167; Mem0.1: 9.0±1.3 vs. 5.5±0.5, t =2.492, df = 11, p = 0.030; Mem1.0: 8.7±1.2 vs. 5.7±1.0, t =2.737, df =10, p = 0.021). Thus, 0.1 and 1.0 mg doses of memantine reversed the age-related recognition memory deficit in rats. However, the effect on DI was not significant after any of the memantine monotreatments (0.1-1.0 mg/kg; F(3, 32) = 0.384; p = 0.765) (Fig. 2B).

**Figure 2.**
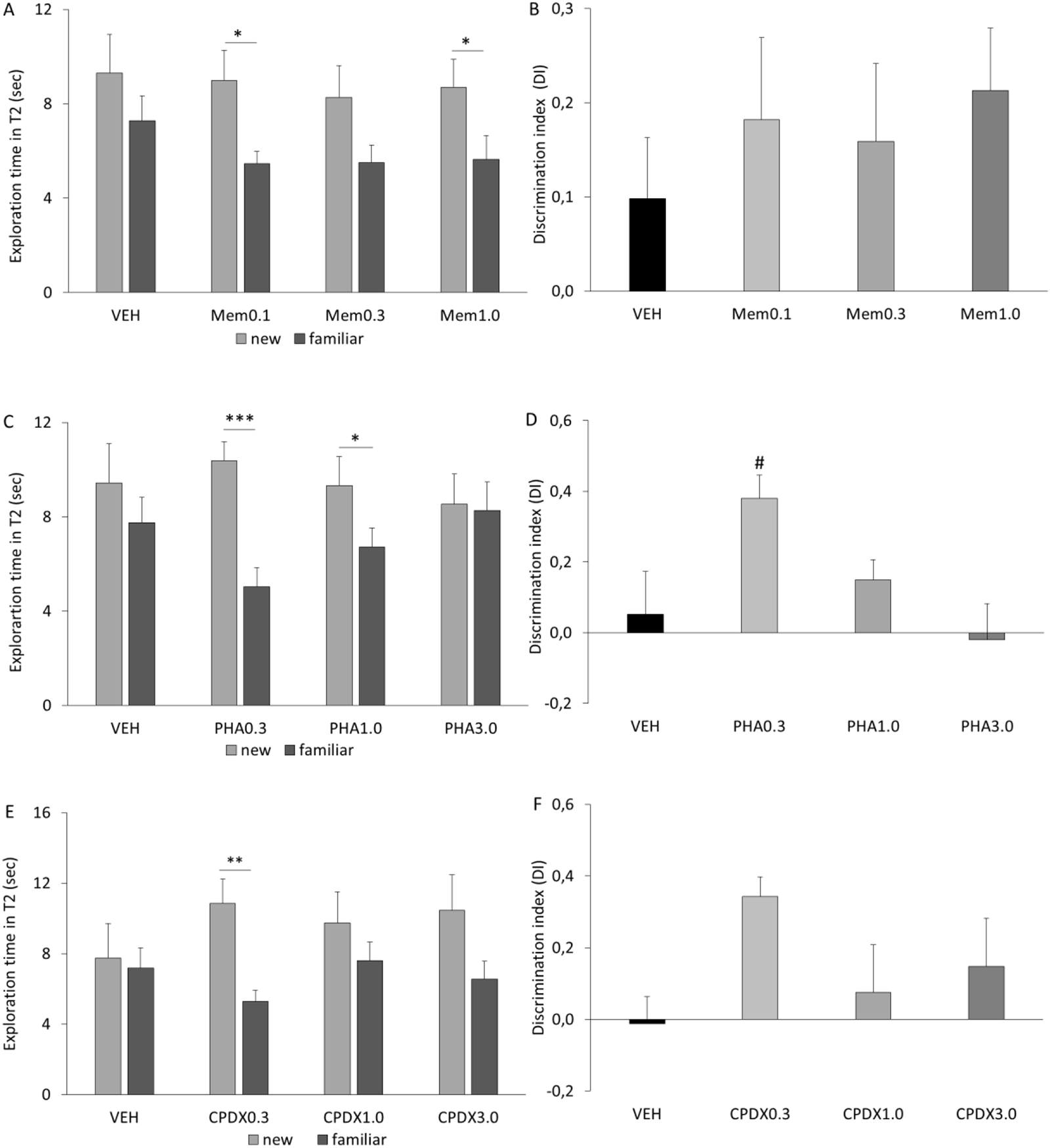
Effects of monotreatments with memantine, PHA-543613 and CompoundX on NOR performance of aged rats. A: Animals received memantine in the lowest (0.1 mg/kg) and in the highest (1.0 mg/kg) dose showed preference towards the exploration of the novel object compared to the familiar one. B: However, DI was not significantly increased by memantine. C: Aged rats that received PHA-543613 in 0.3 mg/kg and 1.0 mg/kg dose explored the novel object for significantly longer time compared to the exploration of the familiar one. D: PHA-543613 in the lowest dose (0.3 mg/kg) improved the DI of aged rats compared to the corresponding vehicle treatment. E: Aged rats that received CompoundX in the lowest dose (0.3 mg/kg) discriminated between the novel and the familiar objects. F: Discrimination index was not significantly increased by CompoundX. Asterisks mark significant difference in the time spent with the exploration of the novel and the familiar objects: ***p<0.001, **p<0.01, *p<0.05 (paired samples T-test). Hash indicates significantly different DI compared to the corresponding vehicle treatment: #p<0.05 (linear mixed-effect model+post-hoc LSD) .

Aged animals receiving PHA-543613 in 0.3 mg/kg and 1.0 mg/kg doses showed a preference towards the novel object (novel vs. familiar: PHA0.3: 10.4±0.8 vs. 5.0±0.8, t =6.354, df = 12, p < 0.001; PHA1.0: 9.3±1.2 vs. 6.7±0.8, t =2.684, df = 12, p = 0.020), while vehicle treated aged rats did not discriminate between the objects (novel vs. familiar: VEH: 9.4±1.7 vs. 7.8±1.1, t=0.799, df=10, p=0.443) (Fig. 2C). Furthermore, PHA-543613 at the lowest dose (0.3 mg/kg) improved the DI of the aged animals compared to the vehicle-treatment (F(3, 34.7) =4.189; p =0.012; PHA0.3 vs. VEH: 0.38±0.07 vs. 0.05±0.12, p =0.012) (Fig. 2D). CompoundX also enhanced memory performance of rats by restoring their ability to discriminate between the novel and familiar objects at 0.3 mg/kg dose (novel vs. familiar: CPDX0.3: 10.9±1.4 vs. 5.3±0.6, t =5.245, df = 9, p = 0.001) (Fig. 2E). Significant main effect of CompoundX monotreatments was not observed on the DI (F(3, 26.6) =1.093; p = 0.369). However, as Fig. 2F shows, the lowest dose of CompoundX (0.3 mg/kg) increased the mean DI similar to the lowest dose of PHA-543613 (0.3 mg/kg).

### 3.3 Co-administration of memantine with α7 nAChR compounds

In the next series of experiments, a subeffective dose of memantine (0.01 mg/kg) was co-administered with subeffective doses of α7 nAChR compounds PHA-543613 (0.1 mg/kg) or CompoundX (0.1 mg/kg). First, the cognitive enhancing effect of memantine-PHA-543613 combination was compared to the effects of the corresponding monotreatments. Results showed that rats after the treatment with memantine alone and in combination with PHA-543613 spent more time exploring the novel object than the familiar object (novel vs. familiar: Mem0.01: 8.1±0.8 vs. 5.4±0.5, t =2.549, df = 10, p = 0.029; Mem0.01&PHA01: 10.1±1.2 vs. 5.0±0.6, t =3.544, df = 11, p = 0.005) in contrast with vehicle treatment (novel vs. familiar: VEH: 6.5±1.0 vs. 7.2±1.1, t =-0.465, df = 10, p =0.652). However, the monotreatment with PHA-543613 was not found effective (novel vs. familiar: PHA0.1: 8.2±1.3 vs. 6.7±1.1, t =1.604, df = 10, p = 0.140) (Fig. 3A). The applied treatments resulted in a marginal effect on DI (F(3, 30) =2.727; p = 0.062). As expected, pairwise comparisons did not show any significant increase of DI by the low-dose monotreatments with memantine and PHA-543613 compared to the vehicle-treated aged control group (Mem0.01 vs. VEH: 0.19±0.07 vs. -0.06±0.12, p=0.081; PHA01 vs. VEH: 0.09±0.07 vs. -0.06±0.12, p= 0.298). In contrast, DI was significantly increased by the combination treatment (Mem0.01&PHA01 vs. VEH: 0.31±0.10 vs. -0.06±0.12, p=0.01) indicating that the combination of the two distinct mechanisms passed a threshold to successfully reverse age-related recognition memory decline (Fig. 3B).

**Figure 3.**
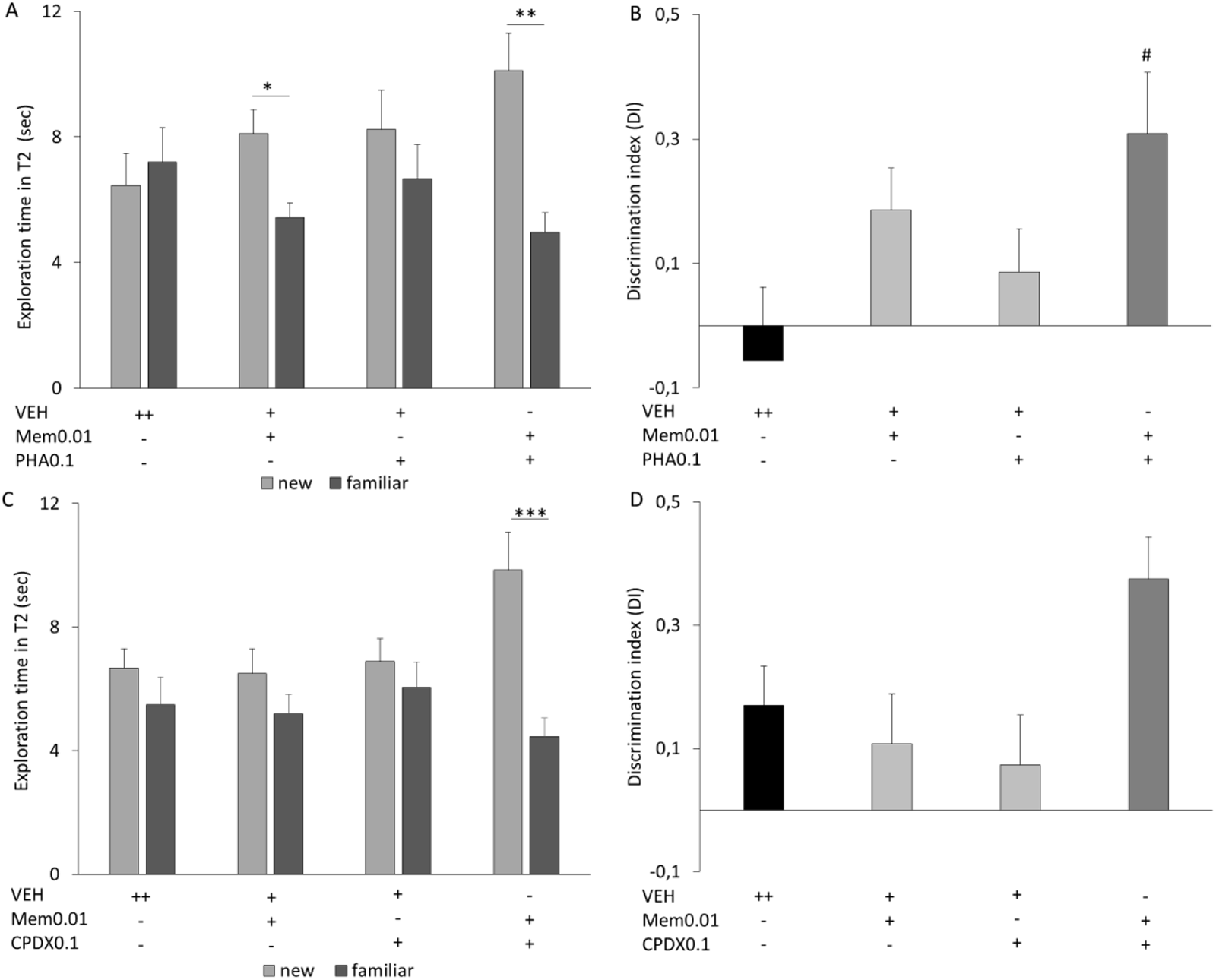
Effects of combination treatments with memantine and α7-nAChR compounds on NOR performance of aged rats. A: Aged rats received memantine alone and in combination with PHA-543613 spent more time exploring the novel object than the familiar one. B: Memantine+PHA-543613 combination treatment significantly improved the DI of aged rats compared to the vehicle-treated group. C: Aged rats that received memantine+CompoundX combination treatment discriminated between the novel and the familiar objects. D: Memantine+CompoundX combination treatment significantly increased the DI of aged rats compared to the vehicle treatment. Asterisks mark significant difference between exploration time of the novel and the familiar objects: ***p<0.001, **p<0.01, *p<0.05 (paired samples T-test). Hash indicates significantly different DI compared to the corresponding vehicle treatment: #p<0.05 (linear mixed-effect model+post hoc LSD) .

Similarly, the effects of the co-administration of memantine and CompoundX were also tested. As expected, animals who received low-dose memantine or CompoundX in monotreatments did not discriminate between the novel and the familiar objects (novel vs. familiar: Mem0.01: 6.5±0.8 vs. 5.2±0.6, t =1.716, df =14, p =0.108; CPDX0.1: 6.9±0.7 vs. 6.1±0.8, t =0.965, df = 13, p = 0.352). However, co-administration of memantine and CompoundX reversed age-related memory deficit as rats spent significantly more time with the exploration of the novel object after the combination treatment (novel vs. familiar: 9.8±1.2 vs. 4.5±0.6, t =4.560, df = 14, p< 0.001) (Fig. 3C). In addition, the combination of memantine and CompoundX also reached a significant effect on DI (F(3, 41.7) =3.281; p =0.030). Regarding monotherapies, age-related decrease of DI was not significantly attenuated by either memantine or CompoundX monotreatments (Mem0.01 vs. VEH: 0.11±0.08 vs. 0.17±0.06, p=0.565; CPDX0.1 vs. VEH: 0.07±0.08 vs. 0.17±0.06, p=0.378). However, the mean DI was marginally significantly increased by memantine-CompoundX co-administration (Mem0.01&PHA01 vs. VEH: 0.37±0.07 vs. 0.17±0.06, p=0.055) (Fig. 3D).

### 3.4. Effects of aging on mRNA expression levels of inflammatory factors and α7 nAChR in the brain

For further biochemical investigations, 5 young and 10 aged animals were sacrificed. Aged animals were further divided into memory-impaired (AI) and unimpaired (AU) aged groups, with 5 animals in each group, depending on their baseline cognitive performance. Rats who performed above the median DI were considered AU (DI range: 0.17 – 0.68), while rats with a DI lower than the median were considered AI rats (DI range: –0.66 – –0.01).

To evaluate the inflammatory profile and cholinergic aging, mRNA expression levels of various cytokines and α7 nAChR were assessed in neocortical, striatal, and hippocampal brain tissue samples of young, AI and AU rats.

Results of qRT-PCR analysis revealed that both AI and AU animals showed significantly higher IL-1β mRNA levels in the neocortex and the striatum compared to young control rats (CTX: F(2, 11) =5.631, p =0.021; AI vs. young: 26.9±4.7 vs. 9.7±2.1, p=0.008; AU vs. young: 22.2±4.5 vs. 9.7±2.1, p=0.047; STR: F(2, 11) =8.753, p =0.005, AI vs. young: 55.3±8.3 vs. 18.9±2.4, p=0.002; AU vs. young: 40.8±7.2 vs. 18.9±2.4, p=0.038) (Fig. 4A-B). On the contrary, only AI animals exhibited a significant increase of IL-1β mRNA in the hippocampus compared to young animals (F(2, 12) =6.708, p =0.013; AI vs. young: 35.6±3.1 vs. 19.1±3.6, p=0.005), and a difference between AU and young groups was not found (AU vs. Young: 23.7±3.4 vs 19.1±3.6, p=0.349) (Fig. 4C).

**Figure 4.**
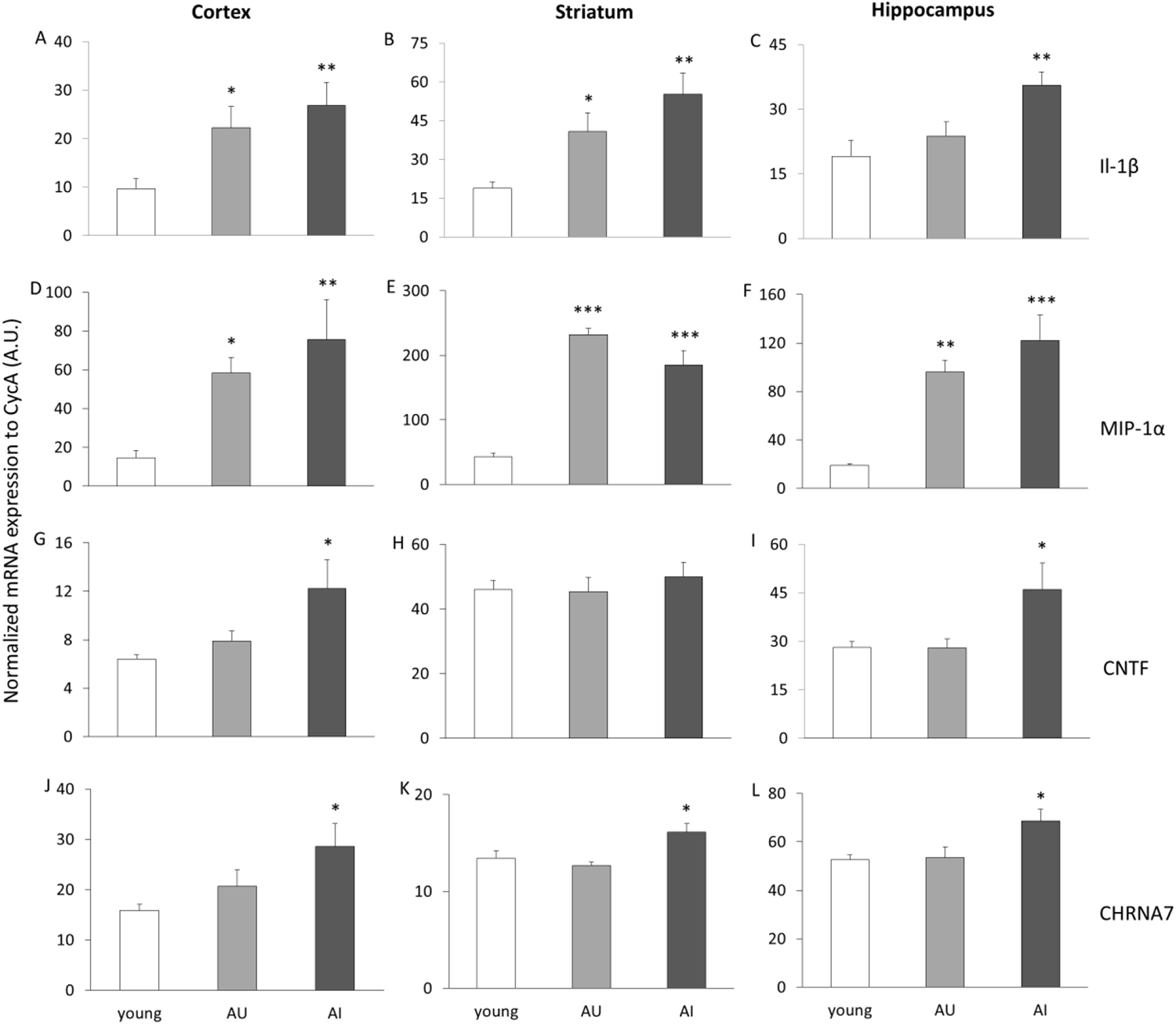
Brain mRNA expression levels of inflammatory markers, CNTF and α7-nAChRs in young, aged cognitively unimpaired (AU) and aged impaired (AI) rats. A-C: Cortical and striatal IL-1β mRNA level significantly increased in both AU and AI groups compared to the young group, while in the hippocampus, only the AI group showed elevated IL-1β mRNA expression. D-F: Significant MIP-1α mRNA upregulation was observed in both AU and AI groups compared to young group in all examined brain areas. G-I: Cortical and hippocampal CNTF mRNA level significantly increased in AI group compared to the young group. J-L: AI animals exhibited α7-nAChR mRNA upregulation in all examined brain areas. Asterisks mark significant changes in mRNA expression levels compared to young animals: ***p<0.001, **p<0.01, *p<0.05 (univariate ANOVA+post-hoc LSD)

MIP-1α mRNA expression was upregulated in both AI and AU rats in all selected brain areas compared to young animals (CTX: F(2, 12) =5.860, p=0.017; AI vs. young: 75.6±20.7 vs. 14.5±3.8, p=0.006; AU vs. young: 58.3±8.0 vs. 14.5±3.8, p=0.035; STR: F(2, 12) =45.5, p <0.001; AI vs. young: 184.8±22.7 vs. 42.6±5.8, p<0.001; AU vs. young: 232.1±9.7 vs. 42.6±5.8, p<0.001; HC: F(2, 12) =16.121, p<0.001; AI vs. young: 122.1±21.1 vs. 18.8±1.6, p >0.001; AU vs. young: 96.2±9.5 vs. 18.8±1.6, p=0.001) (Fig. 4D-F). In the neocortex and the hippocampus the AI group showed the highest mean expression of MIP-1α mRNA (Fig. 4D,F). CNTF mRNA expression levels were significantly increased in the AI group in the cortex and hippocampus compared to young animals (CTX: F(2, 11) =4.167, p =0.045; AI vs. young: 12.2±2.3 vs. 6.4±0.4, p=0.017; HC: F(2, 12) =4.099, p =0.044; AI vs. young: 46.0±8.2 vs. 28.1±1.7, p=0.029). In contrast, AU rats did not express increased mRNA levels of CNTF in these brain regions (CTX: AU vs. young: 7.9±0.8 vs. 6.4±0.4, p=0.507; HC: AU vs. young: 28.0±2.8 vs. 28.1±1.7, p=0.990) (Fig. 4G,I). CNTF mRNA expression in the striatum did not differ between the groups (F(2, 12) =0.394, p =0.683) (Fig. 4H).

AI rats also showed upregulated α7 nAChR mRNA levels in the examined brain areas compared to the young group, while the AU group expressed α7 nAChR mRNA levels similar to the young rats (CTX: F(2, 11) =3.739, p =0.058; AI vs. young: 28.6±4.6 vs. 15.8±1.3, p=0.020; AU vs. young: 20.7±3.2 vs. 15.8±1.3, p=0.350; STR: F(2, 11) =5.635, p =0.021; AI vs. young: 16.1±0.9 vs. 13.5±0.7, p=0.027; AU vs. young: 12.7±0.4 vs. 13.5±0.7, p=0.489; HC: F(2, 12) =3.085, p =0.083; AI vs. young: 68.7±3.0 vs. 52.6±5.4, p=0.047; AU vs. young: 53.4±6.5 vs. 52.6±5.4, p=0.907) (Fig. 4J-L).

### 3.5. Effects of aging on protein expression levels of inflammatory factors and α7 nAChR in the brain

Next, we analyzed the protein expression levels of IL-1β, CNTF, MIP-1α, and α7 nAChR in the neocortex, striatum, and hippocampus of young, AI, and AU rats.

Results of ELISA analysis revealed a significant increase of IL-1β protein level in the striatum of AU but not of AI animals compared to the young control group (F(2, 12) =3.127, p =0.081; AU vs. young: 1.5±0.1 pg/µg vs. 1.1±0.1 pg/µg, p=0.032; AI vs. young: 1.4±0.1 pg/µg vs. 1.1±0.1 pg/µg, p=0.110) (Fig. 5B). In the neocortex and hippocampus no differences in IL-1β expression were found between the groups (CTX: F(2, 7) =0.511, p =0.621; HC: F(2, 12) =0.061, p =0.941) (Fig. 5A,C).

**Figure 5.**
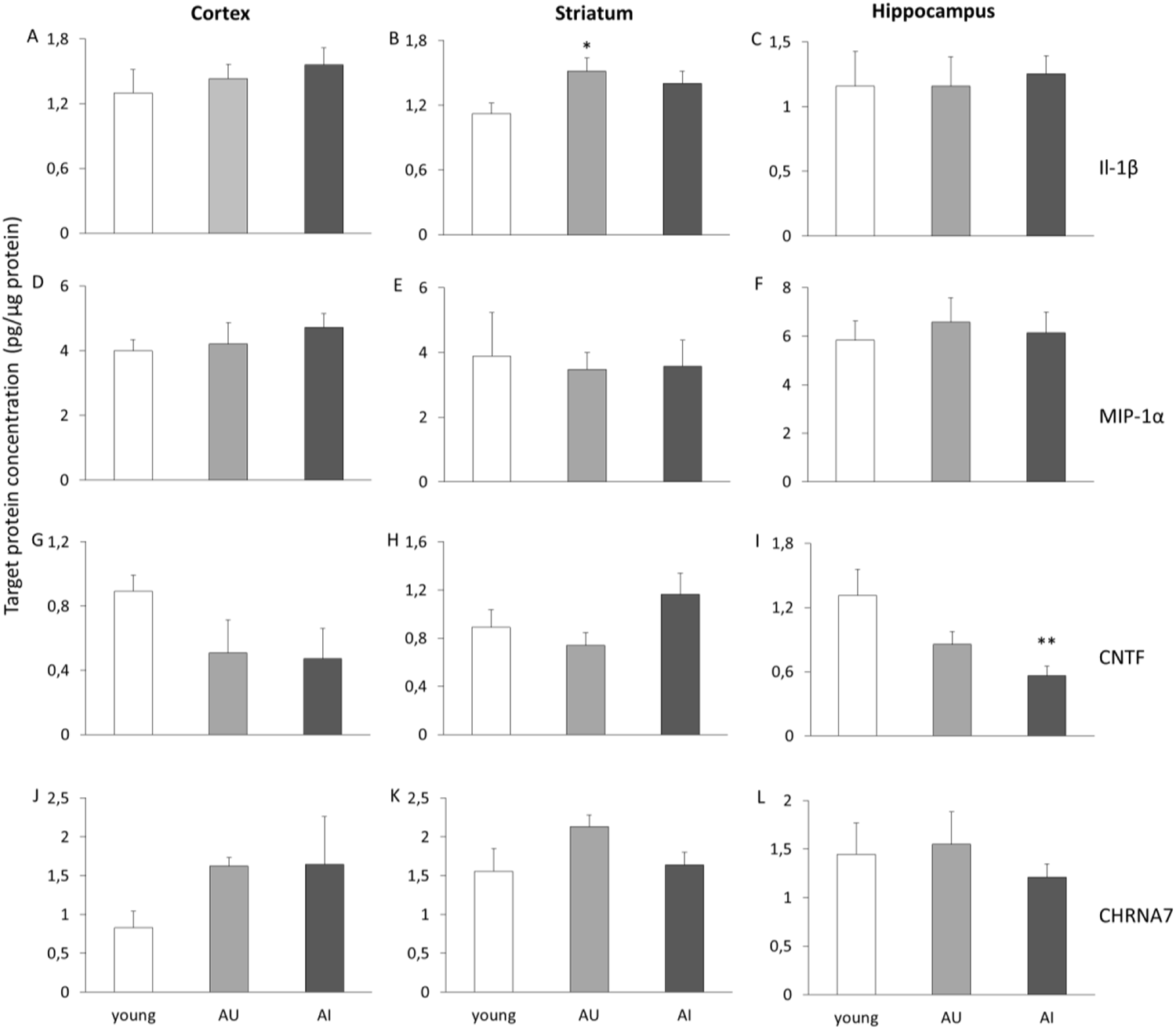
Brain protein expression levels (pg/µg protein) of inflammatory markers, CNTF and α7-nAChRs in young, aged cognitively unimpaired (AU) and aged impaired (AI) rats. A-C: Striatal IL-1β protein level significantly increased in AU group compared to the young group. D-F: MIP-1α protein expression was not different between the groups in any of the tested brain regions. G-I: Hippocampal CNTF protein levels significantly decreased in the AI group compared to the young group. J-L: α7-nAChR protein expression levels were not different between the groups in any of the examined brain areas. Asterisks mark significant changes in protein expression levels compared to young animals: **p<0.01, *p<0.05 (univariate ANOVA+post-hoc LSD).

Surprisingly, neither the AI nor AU groups showed a significant change in the expression of MIP-1α protein in any of the tested brain regions (CTX: F(2, 7) =0.652, p =0.550; STR: F(2, 12) =0.051, p =0.950; HC: F(2, 12) =0.173, p =0.843) (Fig. 5D-F).

In contrast with mRNA expression levels, protein levels of hippocampal CNTF were significantly downregulated in the AI group compared to the young group (F(2, 12) =5.386, p =0.021; AI vs. young: 0.6±0.1 pg/µg vs. 1.3±0.2 pg/µg, p=0.007). In addition, there was a (non-significant) tendency to decreased CNTF protein expression level in the hippocampus of the AU group compared to young animals (AU vs. young: 0.9±0.1 pg/µg vs. 1.3±0.2 pg/µg, p=0.070) (Fig. 5I). In the cortex and the striatum, no significant main effect of aging could be observed on CNTF protein levels of rats (CTX: F(2, 7) =2.298, p =0.171; STR: F(2, 12) =2.167, p =0.151) (Fig. 5G,H).

There was no significant main effect on the protein expression level of α7 nAChR (CTX: F(2, 7) =1.882, p =0.222; STR: F(2, 12) =2.120, p =0.163; HC: F(2, 12) =0.383, p =0.690) (Fig.J-L). However, in the striatum, a (non-significant) tendency of upregulation of α7 nAChR was detected in the AU group compared to the young group (AU vs. Young: 2.1±0.1 pg/µg vs. 1.6±0.2 pg/µg, p=0.081) (Fig. 5K).

## 4. Discussion

The present study provides evidence for the beneficial procognitive effects of combination treatments by co-targeting glutamatergic and cholinergic neurotransmission in the brain using memantine and two α7 nAChR-selective agents in naturally aged rats. We confirmed that aged rats display substantial cognitive impairment compared to young rats which is in line with several earlier preclinical studies (Burger et al. 2007; Long et al. 2009; Santín-Márquez et al. 2021). Monotreatments with memantine or two α7 nAChR ligands could alleviate the age-related cognitive impairment of rats at certain doses. Results showed that memantine at both the lowest (0.1 mg/kg) and highest (1.0 mg/kg) doses was able to improve recognition memory performance of aged rats, while both PHA-543613 and CompoundX exerted their maximum efficacy at the lowest 0.3 mg/kg dose following the usually observed pattern of cholinergic modulation which resembles an inverted U-shaped function (Bentley et al. 2011). Although 5 mg/kg memantine dose can be considered therapeutically relevant for rats (Parsons et al. 2007), there is supporting evidence for the effectiveness of memantine at 1.0 mg/kg or lower doses without undesirable behavioral side-effects. For example, Wise and Lichtman (2007) demonstrated that memantine below the 1.0 mg/kg dose improves the spatial memory of rats in the radial arm maze. In line with that, our latest study also confirmed the effectiveness of memantine at 1.0 mg/kg dose on the long-term spatial memory performance of rats in the Morris water maze task (Bruszt et al. 2021). Moreover, we have earlier reported that memantine, administered in a fairly low dose (0.1 mg/kg) also effectively reversed scopolamine-induced transient short-term memory deficit of rats in the spontaneous alternation task (Bali et al. 2019a). In the same experiment, we found the effective dose of PHA-543613 at 3 mg/kg (Bali et al. 2015). Surprisingly, in the present experiments a 10-fold lower dose of PHA-543613 (0.3 mg/kg) already successfully ameliorated the natural cognitive deficits of aged rats. The difference between the optimal doses of PHA-543613 in the different studies may be explained by the differences in the applied cognitive impairment model, and it suggests that more relevant preclinical data might be acquired using natural models of cognitive impairment such as aged animals in the present study. Other selective α7 nAChR agonists, e.g., AZD0328, were also reported as effective in improving NOR performance of mice at ultra-low doses (0.00178-1.78 mg/kg) (Sydserff et al. 2009; Werkheiser et al. 2011).

Next, we aimed to investigate the effects of co-administration of subeffective doses of memantine (0.01 mg/kg) and PHA-543613 (0.1 mg/kg), or memantine (0.01 mg/kg) and CompoundX (0.1 mg/kg). We found that the long-term recognition memory impairment in aged rats was effectively reserved by memantine-PHA-543613 combination treatment showing a superior cognitive enhancement effect over the corresponding monotreatments. In line with this, the memantine-CompoundX combination treatment also showed remarkable effectiveness, while low-dose monotreatments failed to enhance the recognition memory performance of aged rats. Taken together, the present results indicate a positive interaction between memantine and α7 nAChR compounds, as they reversed age-derived cognitive decline in rats and exceeded the cognitive behavioral effect of the monotherapies. These findings suggest the role of α7 nAChRs in the observed interaction between memantine and α7 nAChR compounds which has been previously corroborated by several preclinical studies. For example, Nikiforuk et al. (2016) demonstrated that higher cognitive flexibility and enhanced recognition memory were observed in rats when memantine was combined with galantamine or selective α7 nAChR PAMs (CCMI or PNU-120596). In the same study, the effects of combination treatments were successfully blocked by the α7 nAChR antagonist methyllycaconitine (MLA), indicating that cognitive enhancement by combination treatment was an α7 nAChR-dependent process. These results are further supported by the study of Busquet et al. (2012), showing that spontaneous alternation and object recognition of mice were facilitated by combining subeffective doses of galantamine and memantine. In addition, we have previously demonstrated that improvement of spatial short-term (working) and long-term memory may also be achieved with combination treatments using the α7 nAChR agonist PHA-543613 and memantine (Bali et al. 2019a; Bruszt et al. 2021).

The exact mechanism underlying the beneficial interaction between α7 nAChR compounds and memantine has not been clarified, yet. In general, according to our present knowledge, exogenous agonists induce the direct, but transient activation of α7 nAChRs, while PAMs rather potentiate the effect of the endogenous ligand (acetylcholine) through positive allosteric modulation of the receptor in addition to reduce its desensitization (Wallace and Bertrand 2015). Positive allosteric modulators seem to be viable therapeutic approaches in NCDs, and α7 nAChR-related mechanisms were determined beneficial in combination treatments as they could similarly facilitate the efficacy of memantine as a cognitive enhancer. An in vitro study in mice brain slices showed that memantine exerts an antagonistic effect on α7 nAChRs and that it shows higher affinity to α7 nAChRs than to NMDA receptors (Aracava et al. 2005). In addition, α7 nAChR antagonist methyllycaconitine (MLA) has been proven to improve memory acquisition in rats at low doses, suggesting a final cognitive effect similar to that of the agonists of the α7 nAChR (van Goethem et al. 2019). Accordingly, it can be concluded that the action of memantine on α7 nAChRs may also be involved in the pro-cognitive effects of the combination treatments, especially at low doses, thus α7 nAChRs may serve as the common targets of memantine and PHA-543613 or CompoundX. Our results clearly demonstrated that two distinct mechanisms of action – activation of nAChRs and memantine-induced multiple cellular effects on different neurotransmitter receptors – can result in a synergistic action to alleviate the observed age-related decline in cognitive performance.

Furthermore, in our current study, mRNA and protein expression levels of multiple immunological markers have been investigated, because increased state of neuroinflammation in the aging brain can be considered main pathological hallmark of NCDs (Ownby 2010; Ahmad et al. 2022). To evaluate the differences between normal and pathological aging on the cellular level, the neuroinflammatory state of aged rats with normal and impaired memory performance was compared. We demonstrated a similar pattern of mRNA expression levels of IL-1β and MIP-1α in almost all examined brain areas between the groups. Results indicate that both AU and AI groups exhibited an elevated mRNA upregulation of IL-1β and MIP-1α, which was generally more pronounced during ‘pathological’ aging suggesting a shift toward a pro-inflammatory state of the brain of AI animals. However, the increased mRNA expression was manifested in higher protein levels only in the case of IL-1β in the striatum of AU rats compared to young rats. These data are in agreement with a recent study assessing the protein expression of a wide range of pro-inflammatory markers (Perkins et al. 2021). They found no evidence of elevated expression of most basal cytokines and chemokines using total tissue content analysis in aged animals.

Neurotrophic factors such as BDNF and CNTF are known to be neuroprotective since they can promote synaptic plasticity and neuronal survival (Hagg et al. 1992; Garcia et al. 2010; Askvig and Watt 2019; Wu et al. 2020). Indeed, in our study, CNTF protein expression was significantly lower in the hippocampus of aged animals with memory impairment compared to young rats, and a tendency was also observed towards lower protein levels in the cortex of aged rats. Surprisingly, the cortical and hippocampal CNTF mRNA was upregulated in AI groups. The opposite direction of change in the mRNA and protein levels might be explained by a mechanism that aims at the compensation of lower neurotrophic support at the transcriptional level during aging. However, the reason why the upregulation of CNTF mRNA expression may not be manifested in increased protein expression levels will be investigated by future studies. Aged animals may serve as useful models of NCDs not just because of the elevated neuroinflammation in the brain but also the cholinergic deficits in NCDs, that are accounted for contributing to the decline of learning and memory processes. Therefore, the expression of α7 nAChR mRNA and protein was also investigated in AI and AU rats. We found that AI rats exhibited a robust upregulation of α7 nAChR mRNA in all examined brain regions with no significant changes in the α7 nAChR expression at the protein level. However, it must be noted that a tendency toward increased mean protein expression of α7 nAChR was observed in the neocortex. Probably, underestimated sample sizes and the low sensitivity of the ELISA method compared to RT-PCR may explain why no systematic increase in protein levels could be statistically confirmed as a result of the upregulation of α7 nAChR mRNA in the brain of aged rats. Previous body of evidence in the literature is also ambiguous on the change of α7 nAChR expression in the brain during aging. Several studies found that expression of α7 nAChRs markedly decreased in NCDs, mainly in brain areas related to cognitive functions such as the frontal cortex and hippocampus (Burghaus et al. 2000; Wevers et al. 2000; Hernandez et al. 2010; Ren et al. 2020). However, opposite findings were also reported for age-related upregulation of α7 nAChRs, particularly on the mRNA level, which was also confirmed by our current observations (Nordberg 2001; Kihara and Shimohama 2004; Conejero-Goldberg et al. 2008). Nevertheless, a supposed upregulation of α7 nAChRs may also explain the fairly low effective doses of selective α7 nAChR agents in aged rats, namely that PHA-543613 ameliorated the cognitive deficit of aged rats in a 10-fold lower dose in the present study compared to that in scopolamine-treated young rats in our earlier studies (Bali et al., 2015, 2019a).

In conclusion, in the present study, we confirmed the beneficial interaction between memantine and α7 nAChR acting agents in naturally aged rats demonstrating a translationally more relevant cognitive improvement that might be a target for future procognitive therapy. The relevance of naturally aged rats for modeling human NCDs was also demonstrated by the animals showing characteristic neuropathological changes also at the molecular level, such as elevated levels of neuroinflammatory markers and decreased levels of neurotrophic support to the cells. Combination treatment with memantine and PHA-543613 or memantine and CompoundX successfully alleviated the age-related cognitive decline, moreover efficacy of both combinations was superior over the corresponding monotreatments. Based on our results, we further hypothesize a prominent role of α7 nAChR in the cognitive enhancer effects of the combination treatments in clinical conditions with cognitive decline in older ages. Therapeutic potential of combination treatments that are based on pharmacological interactions on the α7 nAChRs is also supported by the finding that expression of the receptors may become highly upregulated in aging, thus, providing well-accessible binding sites for their pharmacological modulation.

## Acknowledgements

Behavioral experiments were performed in collaboration with the Animal Facility at the Szentágothai Research Centre of the University of Pécs. We are grateful to Kitti Schmeltzer for animal care.

The scientific work/research and/or results disclosed in this article were reached with the sponsorship of the Gedeon Richter Talentum Foundation in the framework of the Gedeon Richter Excellence PhD Scholarship of Gedeon Richter Plc.

## 5. Funding

This study was supported by the UNKP-22-3 new National Excellence Program of the Ministry for Culture and Innovation from the source of the National Research, Development and Innovation Fund (NB) and by the National Research, Development and Innovation Office of the Hungarian Government (grant no. ‘K 129247’, IH). The funders had no role in the design of the study, in the collection, analysis, or interpretation of data, in the writing of the manuscript, or in the decision to publish the results.

## 6. Author Contributions

NB: Methodology, Investigation, Visualization, Formal analysis, Data curation, Writing; ZKB: Conceptualization, Methodology, Investigation, Formal analysis, Data curation, Writing; LVN: Methodology, Investigation, Data curation, Writing; IL: Conceptualization, Writing; LB: Conceptualization, Writing; ZN: Conceptualization, Writing; IH: Conceptualization, Methodology, Supervision, Writing, Resources, Funding acquisition.

## 7. Data Availability Statement

Data is available on request from the authors.

## 8. Conflicts of Interest

IL, ZN, and BL are employees of Gedeon Richter Plc. This does not alter the authors’ adherence to journal policies on sharing data and materials. The remaining authors are academic researchers and declare that the research was conducted in the absence of any commercial, non-financial or financial relationships that could be construed as a potential conflict of interest.

## Notes

### Competing Interest Statement

IL, ZN, and BL are employees of Gedeon Richter Plc. This does not alter the adherence of the authors to journal policies on sharing data and materials. The remaining authors are academic researchers and declare that the research was conducted in the absence of any commercial, non-financial or financial relationships that could be construed as a potential conflict of interest.

